# Metaplastic neuromodulation via transcranial direct current stimulation has no effect on corticospinal excitability and neuromuscular fatigue

**DOI:** 10.1101/2024.02.25.581971

**Authors:** Madison R Boda, Lavender A Otieno, Ashleigh E Smith, Mitchell R Goldsworthy, Simranjit K Sidhu

**Author notes:** Present Address. Behaviour-Brain-Body Research Centre, Justice and Society, University of South Australia, South Australia, Australia (Mitchell R Goldsworthy). Hopwood Centre for Neurobiology, Lifelong Health Theme, South Australian Health and Medical Research Institute, South Australia, Australia (Mitchell R Goldsworthy). **Correspondence**: Dr Simranjit K Sidhu, S433, Helen Mayo South, Frome Rd The University of Adelaide, South Australia, 5005 AUSTRALIA, T.

## Abstract

Transcranial direct current stimulation (tDCS) is a non-invasive brain stimulation tool with potential for managing fatigue, possibly due to alterations in corticospinal excitability. However, inconsistencies in intra– and inter-individual variability responsiveness to tDCS limit its clinical use. Emerging evidence suggests harnessing homeostatic metaplasticity induced via tDCS may reduce variability and boost its outcomes, yet little is known regarding its influence on fatigue in healthy adults. We explored whether cathodal tDCS (ctDCS) prior to exercise combined with anodal tDCS (atDCS) could augment corticospinal excitability and attenuate fatigue. 15 young healthy adults (6 males, 22 ± 4 years) participated in four pseudo-randomised neuromodulation sessions: sham stimulation prior and during exercise, sham stimulation prior and atDCS during exercise, ctDCS prior and atDCS during exercise, ctDCS prior and sham stimulation during exercise. The exercise constituted an intermittent maximal voluntary contraction (MVC) of the right first dorsal interosseous (FDI) for 10 minutes. Fatigue was quantified as an attenuation in MVC force, while motor evoked potential (MEP) amplitude provided an assessment of corticospinal excitability. MEP amplitude increased during the fatiguing exercise, whilst across time, force decreased. There were no differences in MEP amplitudes or force between neuromodulation sessions. These outcomes highlight the ambiguity of harnessing metaplasticity to ameliorate fatigue in young healthy individuals.

## INTRODUCTION

Neuromuscular fatigue is characterised by an exercise-induced decline in maximal voluntary muscle force (Taylor et al., 2016). It is a prevalent incapacitating symptom recognised as causing decrements in quality of life and disability (Kluger et al., 2013). Both the peripheral and central nervous systems of the body are established to contribute to fatigue (Wan et al., 2017). During fatigue, a gradual decrease in maximal voluntary activation of muscle due to a decline in neural drive occurs centrally (Gandevia, 2001). In contrast, alterations distal or within the neuromuscular junction are apparent with peripheral fatigue (Wan et al., 2017). Prior research has also revealed alterations in the excitability of the corticospinal tract during neuromuscular fatigue (Taylor et al., 1996). In fact, corticospinal excitability is enhanced during single-joint fatiguing exercise, possibly reflecting an increase in motor output to the muscle to counteract the progressive difficulty in maintaining contractile force (Gandevia et al., 1996).

The use of non-invasive brain stimulation techniques such as transcranial direct current stimulation (tDCS) can modify corticospinal excitability (Nitsche and Paulus, 2000). tDCS is a portable, non-invasive, painless method of neuromodulation that when employed, delivers continual, low direct electrical current through electrodes on the scalp (Nitsche and Paulus, 2000). The effects of tDCS are predominantly a consequence of polarity-specific bidirectional alteration of resting membrane potential (Liebetanz et al., 2002). Anodal tDCS (atDCS) increases cerebral excitability by eliciting subthreshold membrane depolarisation (Nitsche and Paulus, 2000). On the other hand, cathodal tDCS (ctDCS) causes a decrease in cerebral excitability and membrane hyperpolarisation (Nitsche and Paulus, 2000). Prolonged use of atDCS and ctDCS induces lasting effects similar to persistent forms of activity-dependent synaptic plasticity, namely, long-term depression (LTD) and long-term potentiation (LTP) respectively (Bliss and Lomo, 1973).

Metaplasticity is referred to regulation of neural activity by which induction of synaptic change is dependent on previous synaptic activity (Abraham and Bear, 1996). It operates to preserve synaptic activity within a dynamic range to aid in the integration of temporally dispersed episodes of synaptic change (Abraham and Bear, 1996). The Bienenstock-Cooper-Munro theory presents a theoretical explanation on metaplasticity, implying that the threshold for synaptic modification dynamically and bi-directionally varies as a function of prior activity (Bienenstock et al., 1982). Previous LTP shifts the modification threshold to the right, making further LTP induction more difficult, and therefore increasing the likelihood of LTD (Abraham and Tate, 1997). The opposite is noticed with previous LTD activity (Abraham and Tate, 1997). Prior studies have used two distinct blocks of tDCS over the motor cortex to harness metaplasticity and improve skill acquisition and motor learning in a healthy population (Christova et al., 2015; Fujiyama et al., 2017). For instance, Christova and colleagues found ctDCS preceding atDCS applied concurrently during a grooved pegboard test reduced the completion time of the task when compared to sham condition (Christova et al., 2015). Ultimately, employing priming ctDCS to reduce the modification threshold appears to enhance both corticospinal excitability and functional outcomes of succeeding atDCS (Christova et al., 2015; Fujiyama et al., 2017). Presently, there is a paucity of studies exploring the interaction between metaplastic neuromodulation and neuromuscular fatigue using tDCS.

Currently, no universally efficacious therapy exists for offsetting neuromuscular fatigue (Wan et al., 2017). Given the partial efficacy of existing treatments and the degree to which neuromuscular fatigue can impact an individual’s activities of daily living, exploring the relationship between metaplastic neuromodulation and neuromuscular fatigue may be important for uncovering a non-pharmacological and non-invasive treatment approach. Specifically, we explored priming ctDCS prior to single-joint fatiguing exercise combined with atDCS on corticospinal excitability and neuromuscular fatigue in young, healthy adults. We hypothesised that ctDCS primed atDCS applied concurrently with fatiguing exercise would augment corticospinal excitability facilitation and attenuate fatigue compared to atDCS primed by sham stimulation (stDCS).

## EXPERIMENTAL PROCEDURES

### Participants

Fifteen young healthy volunteers were recruited for the study (6 males, mean age ± SD, 22 ± 4 years). According to the Edinburgh handedness questionnaire, all but one participant were right-handed (handedness Laterality Index, 0.84 ± 0.3), with one individual being ambidextrous (Oldfield, 1971). Before participation, volunteers were screened for contraindications to transcranial magnetic stimulation (TMS) (e.g., metallic implants in the skull, cardiac pacemaker, neurological condition, substance abuse, history of seizures, epilepsy, and/or pregnancy). Participants were instructed to refrain from consuming caffeine 4 hours prior to the experimental session since ingestion of caffeine has been reported to affect fatigability and performance outcomes (Doherty et al., 2004). The protocol was conducted in accordance with the Declaration of Helsinki and was approved by the Human Research Ethics Committee at the University of Adelaide. Written informed consent was obtained from all participants before involvement. Participants were reimbursed for their time upon completion of the study.

### Experimental set-up

Participants sat with their right elbow fixed approximately 90°, pronated forearm aligned on a horizontal surface, and index finger positioned adjacent a force transducer (MLP 100; Transducer Techniques, Temecula, CA). A custom manipulandum was used to restrain the forearm and wrist. Responses evoked from the right first dorsal interosseous (FDI) muscle were recorded via two electromyography (EMG) surface electrodes (Ag/AgCl) arranged over the muscle in a belly-tendon montage. An additional two grounding electrodes were placed on the wrist and forearm to minimise electrical noise. EMG was amplified (1000x) and band-pass filtered (20 Hz – 1 kHz) (1902; Cambridge Electronic Design [CED], UK) prior to digitisation via a 1401 interface at 2 kHz and stored offline.

### Experimental protocol

Participants attended the laboratory for four experimental sessions, each separated by at least 7 days to avoid any long-term effects of tDCS (Peters et al., 2013; Reis et al., 2009). To control for diurnal influence, experimental sessions were completed in the afternoon (after 12 pm) (Ridding and Ziemann, 2010). During each session, the tDCS paradigm was changed and the order was counterbalanced. The order of these sessions was pseudorandomised and double-blinded with the aid of a separate experimenter, meaning both the participant and main experimenter were blinded to tDCS polarities. Within each of the four experimental sessions, tDCS was applied twice. The initial tDCS functioned to prime subsequent tDCS which was applied during exercise. Priming neuromodulation was either stDCS or ctDCS, whilst stDCS or atDCS was applied during the exercise. Hence, the four experimental sessions were: 1) stDCS-stDCS, 2) stDCS-atDCS, 3) ctDCS-atDCS, 4) ctDCS-stDCS. The stDCS-stDCS and ctDCS-stDCS sessions were control conditions, providing evidence on whether the effects of ctDCS-atDCS were explicitly related to the metaplastic neuromodulation intervention, or merely a result of the fatiguing exercise task.

Before employing tDCS, baseline measures of force and corticospinal excitability were taken (Fig. 1(A)). Maximal voluntary contraction (MVC) force was established by calculating the average force of three brief (∼5-second) maximal FDI abduction tasks, each separated by 30-seconds. Baseline corticospinal excitability was measured via eliciting 15 single TMS pulses and 3 peripheral nerve stimulations (PNS). Participants then received priming tDCS at rest; ctDCS or stDCS. At 2 minutes and 8 minutes post-priming, measurements (15 TMS and 3 PNS) were taken to assess the influence of priming on corticospinal excitability. 10 minutes succeeding the termination of priming, test tDCS stimulation begun; atDCS or stDCS (Fig. 1(B)). An inter-stimulation interval of 10 minutes deemed suitable to ensure test tDCS was executed throughout the after-effects of priming tDCS (Monte-Silva et al., 2010). At the start, middle and end of all tDCS periods, participants reported sensations from the tDCS electrodes. 30-seconds following test tDCS commencement, corticospinal excitability measurements (15 TMS and 3 PNS) were repeated. Participants then completed the fatiguing exercise comprising 10 intermittent 30-second MVCs. Between each set (30-seconds), 5 TMS and 1 PNS were elicited. Participants were verbally instructed to start and stop contracting as well as verbally encouraged to perform maximally during the exercise. Force and EMG output were presented on a computer monitor for visual feedback. Post-exercise measurements (15 TMS, 3 PNS and 2 brief MVCs separated by 30-seconds) were completed immediately following exercise and repeated 10 minutes and 20 minutes after the exercise concluded to gauge fatigue recovery and tDCS after-effects (Fig. 1(C)).

**Fig. 1.**
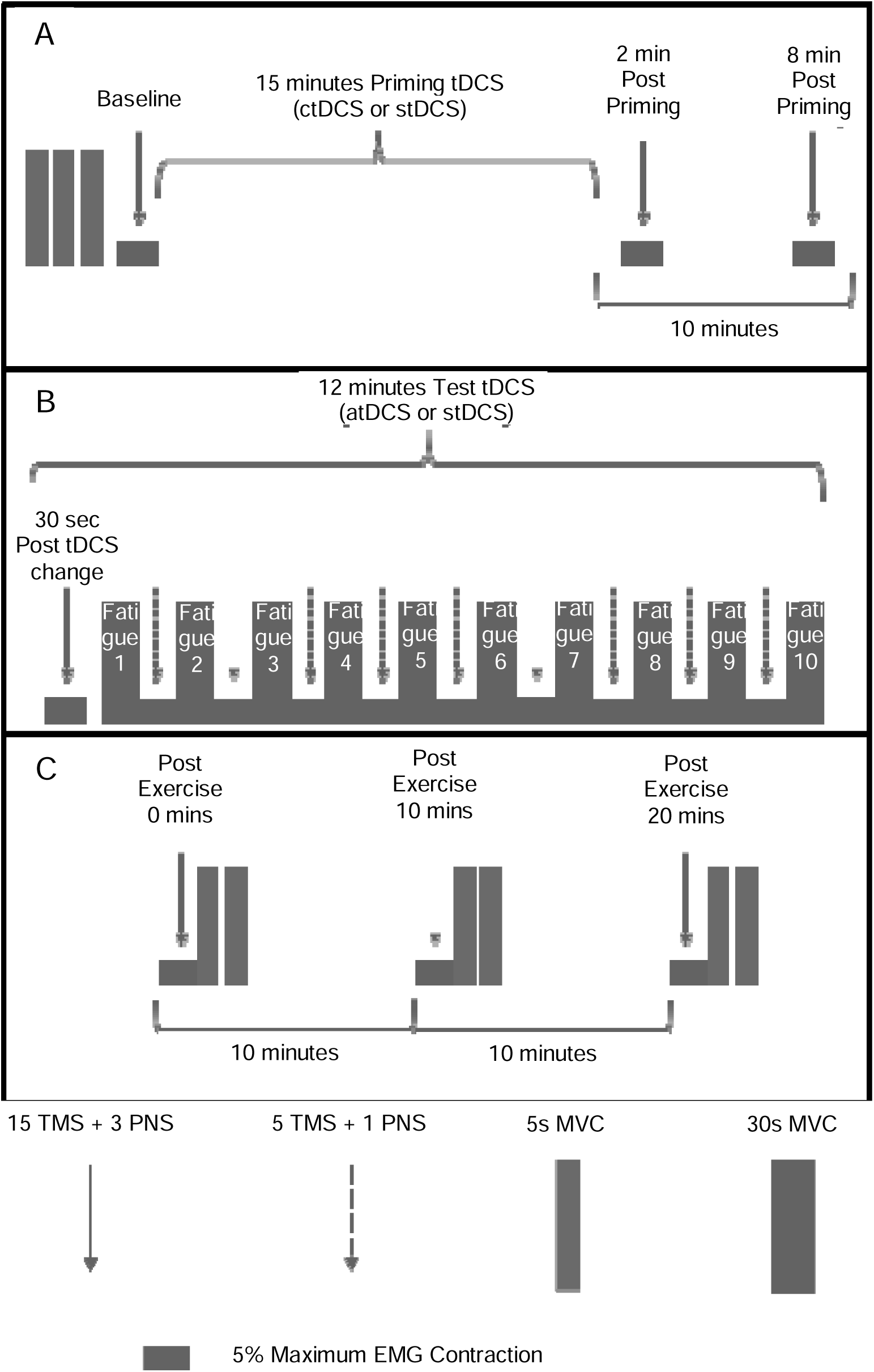
Experimental protocol schematic. (A) Baseline and priming tDCS (B) Fatiguing exercise and test tDCS (C) Post-exercise/recovery

### tDCS

Constant direct current was delivered by a pair of 35 cm^2^ saline-soaked, synthetic sponge electrodes connected to a battery-powered direct current stimulator (NeuroConn DC Stimulator Plus, Germany). A bipolar tDCS electrode montage appears to optimally enhance motor cortex excitability (Nitsche and Paulus, 2000) and hence, was used in the current study. The active electrode was centred on the left primary motor cortex (M1), specifically over the representational field of the right FDI muscle. The reference electrode was positioned over the contralateral supraorbital region. tDCS polarity refers to the electrode over the left M1. The M1 hotspot for tDCS was determined as the area of the left hemisphere exhibiting the largest motor evoked potential (MEP) amplitude at 40-60 % of maximum TMS stimulator output. The current of the tDCS was set at an intensity of 1 mA (Batsikadze et al., 2013; Jamil et al., 2017; Xian et al., 2023). To replicate previous work, priming ctDCS was applied for 15 minutes, whilst atDCS during exercise (test tDCS) was administered for 12 minutes (Fig. 1) (Xian et al., 2023). For stDCS, although the electrode montage was identical to that of genuine tDCS administration, current was only delivered for 30-seconds. Thus, no corticospinal excitability changes were induced during stDCS (Kristiansen et al., 2021). In both real and stDCS, the current was ramped-up and down over 8-seconds at the start and end of the stimulation to inhibit electrical transients.

### TMS

TMS was applied to the left M1 to evaluate the excitability of projections from the cortical representation of the right FDI muscle. Single-pulse TMS stimuli were delivered using a standard figure-of-8 magnetic coil (9 cm external wing diameter) connected to a Magstim BiStim unit (MagStim Company, Dyfed, UK). The coil was positioned tangentially over the brain at a 45° angle to the sagittal plane to deliver a posterior-anterior current flow, and to optimally elicit MEPs in the FDI. The amplitude of MEP was used to assess corticospinal excitability. All TMS pulses were delivered while participants contracted their right FDI muscle at 5 % of their maximum EMG (as determined during baseline MVC trials). An active state was chosen for practicality since a lower stimulation intensity is required to consistently produce a MEP if muscle is active (Ngomo et al., 2012). Additionally, and perhaps more importantly, active muscle more closely represents what occurs during a task (i.e., fatiguing exercise). The FDI motor hotspot was established by mapping (at 40-60 % of maximum stimulator output) for the area that resembled the largest MEP amplitude. The position of the coil was marked on the tDCS electrodes and scalp with permanent marker to aid in consistent coil positioning. These markings were continuously monitored throughout the protocol. Active motor threshold (AMT) was defined as the lowest TMS stimulus intensity necessary to elicit a MEP discernible from background EMG signal in 3 out of 5 trials (Sidhu et al., 2017). TMS intensity was set at 120 % of AMT for all measurement blocks (55 ± 10 % of maximum stimulator output) to ensure effective activation of the FDI during the experiment.

### PNS

The right ulnar nerve was peripherally stimulated via a bipolar bar electrode probe connected to a constant-current stimulator (DS7A; Digitimer, Hertfordshire, UK). Upon establishing the optimal position, the probe was secured so that the cathode was angled distally. The optimal position was established as the site eliciting the greatest compound muscle action potential (M-wave) in resting FDI at 10 mA current. To ascertain the maximum M-wave (Mmax), stimulation intensity was incrementally increased by 5 mA until the M-wave amplitude ceased to rise. Stimuli were delivered at a test intensity of 120 % of the intensity necessary to produce Mmax (27 ± 6.4 mA).

### Data analysis

MVC data were manually analysed using offline recordings on Spike2 software (Version 6.18; CED, UK). Force amplitude was measured during brief (∼5-second) MVCs at baseline and post-exercise and averaged across trials at each time point. Mean force (from initial peak to just prior to the end of contraction) was measured for each of the 30-second MVCs during the exercise. The root mean squared EMG was measured for every 5 % maximum EMG contraction to ensure all measurements were taken at a constant EMG level.

MEP and Mmax amplitudes were calculated in millivolts using offline recordings on Spike2 software. MEP amplitude at individual time points were determined as the mean amplitude across all trials in the measurement block. MEPs were normalised to Mmax (MEP % Mmax) to study corticospinal changes whilst excluding changes at the level of the muscle.

### Statistical analysis

IBM SPSS Statistics software (Version 24; Chicago, USA) was used for the statistical analyses. Separate linear mixed model (LMM) analyses with factors time and neuromodulation (stDCS-stDCS vs. stDCS-atDCS vs. ctDCS-atDCS vs. ctDCS-stDCS) were used to assess main effects and interactions. For MEP, LMM analyses were performed for baseline and post-priming, 30-seconds post-tDCS and all the fatiguing contraction time points, and baseline and post-exercise. Furthermore, two LMM analyses were completed for force: one for all the fatiguing contraction time points and another for baseline and post-exercise. For all comparisons, normality of the data was validated using Shapiro-Wilk tests. Post hoc tests with Bonferroni’s correction for multiple comparisons were employed to probe significant main and interaction effects. One-way analyses of variance were used for evaluating differences between neuromodulation conditions in time of day, lab temperature and humidity. All data in text and tables are conveyed as mean ± SD. Data in figures are illustrated as box-and-whisker plots, with the “box” depicting the median and the 25th and 75th quartiles, and the “whisker” highlighting the 5th and 95th percentile. The significance of outcomes was set at P < 0.05.

## RESULTS

Participant characteristics are displayed in Table 1. No adverse reactions to tDCS were reported by participants. Participants were unable to distinguish stDCS from real tDCS since sensations related with real stimulation were also communicated during stDCS. The sensations reported comprised needling, warmth, burning, stinging, prickling, itchiness, and tingling. Most described sensations at the beginning of tDCS but, felt nothing when asked in the middle and at the end of the stimulation. There were no differences between neuromodulation conditions in time of day (1:26 pm ± 1.2 hrs; P ≥ 0.679), lab temperature (21.6 ± 1.1°C; P ≥ 0.426), or humidity (40.0 ± 5.1 %; P ≥ 0.124). All data were normally distributed with equal variances assumed.

**Table 1:** Demographic characteristics of participants. Work, sport, and leisure time indices are sub-scales of the International Physical Activity Questionnaire (Baecke et al., 1982). Participant scores ranged from 1 (sedentary) to 5 (active) for each index.

| Characteristic | Mean ± SD |
| --- | --- |
| Height (cm) | $170.1 \pm 8.0$ |
| Weight (kg) | $64.5 \pm 12.0$ |
| Handedness | $0.8 \pm 0.3$ |
| Work index | $2.8 \pm 0.9$ |
| Sport index | $2.3 \pm 0.8$ |
| Leisure time index | $2.6 \pm 0.8$ |

### Corticospinal excitability

Post-priming, there were no main effects of time (F_2,10_ = 0.017, *P* = 0.984), neuromodulation (F_3,13_ = 2.00, *P* = 0.163) nor interaction between time and neuromodulation (F_6,87_ = 1.12, *P* = 0.360) on MEP amplitudes (Fig. 2(A)). During exercise (Fig. 2(B)) however, there was a main effect of time (F_9,123_ = 4.82, *P* < 0.001) on MEP amplitudes but, not neuromodulation (F_3,55_ = 1.27, *P* = 0.293) nor interaction between time and neuromodulation (F_27,380_ = 1.02, *P* = 0.444). MEPs were facilitated compared to 30-seconds post-test tDCS across all neuromodulation conditions during fatiguing contraction two through to the ninth contraction (*P* < 0.05). Similarly, there was a main effect of time (F_3,33_ = 4.14, *P* = 0.013) on MEP amplitudes post-exercise (Fig. 2(C)) but, not neuromodulation (F_3,48_ = 0.396, *P* = 0.756) nor interaction between time and neuromodulation (F_9,131_ = 1.67, *P* = 0.103). Across all neuromodulation conditions, MEP increased from baseline immediately post-exercise (0 min) (P = 0.008) but returned to baseline 10 minutes post-exercise (P = 0.923).

**Fig. 2.**
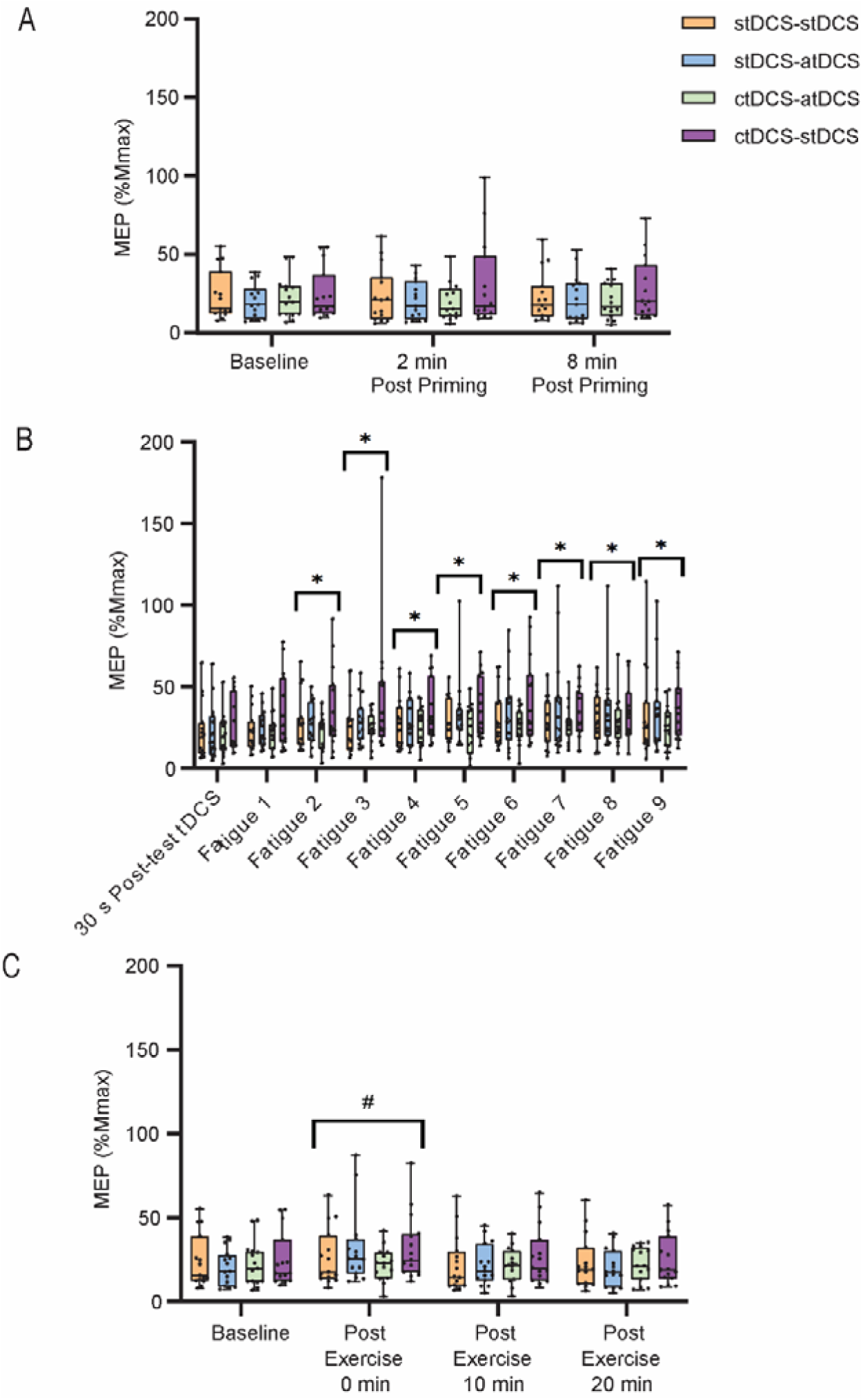
Mean MEP amplitude (expressed as percentage of Mmax) post-priming (A), during fatiguing exercise (B), and post-exercise (C) from fifteen young healthy adults displayed across neuromodulation conditions (stDCS-stDCS, stDCS-atDCS, ctDCS-atDCS, ctDCS-stDCS) over time. Bonferroni’s post hoc tests: * indicates significant difference from 30s post-test tDCS (*P* < 0.05); # denotes significant difference from baseline (*P* < 0.05)

Post-priming, there were no main effects of time (F_2,27_ = 0.532, *P* = 0.593), neuromodulation (F_3,55_ = 1.77 *P* = 0.163) nor any interaction between time and neuromodulation (F_6,85_ = 0.688, *P* = 0.660) on Mmax (baseline Mmax across all neuromodulation conditions, 15.7 ± 0.58 mV). However, during exercise, there was a main effect of time (F_9,126_ = 25.4, *P* < 0.001) on Mmax but, not neuromodulation (F_3,56_ = 1.13, *P* = 0.343). There was also an interaction between time and neuromodulation (F_27,378_ = 1.88, *P* = 0.006) on Mmax during exercise. Mmax attenuated from 30-seconds post-test tDCS across all neuromodulation conditions (*P* ≤ 0.004) during fatiguing contraction one through to the ninth contraction. Additionally, at 30-seconds post-test tDCS, Mmax was greater in stDCS-atDCS (16.6 ± 4.0 mV, *P* = 0.034) and ctDCS-stDCS (16.6 ± 4.0 mV, *P* = 0.036) neuromodulation conditions when compared to stDCS-stDCS (13.3 ± 4.0 mV). Post-exercise, there was a main effect of time (F_3,39_ = 21.9, *P* < 0.001) on Mmax but, not neuromodulation (F_3,53_ = 1.40, *P* = 0.252) nor interaction between time and neuromodulation (F_9,127_ = 0.91, *P* = 0.519). Across all neuromodulation conditions, Mmax attenuated from baseline immediately post-exercise (0 min) (*P* < 0.05) but returned to baseline from 10 minutes post-exercise (*P* ≥ 0.218).

There were no main effects of time (F_2,27_ = 0.745, *P* = 0.484), neuromodulation (F_3,55_ = 1.01 *P* = 0.395), nor any interaction between time and neuromodulation (F_6,84_ = 1.74, *P* = 0.121) on background EMG post-priming. Similarly, there were no main effects of time (F_9,504_ = 0.761, *P* = 0.652), neuromodulation (F_3,56_ = 1.01 *P* = 0.397), nor any interaction between time and neuromodulation (F_27,504_ = 1.20, *P* = 0.225) on background EMG during exercise. There were also no main effects of time (F_3,42_ = 0.305, *P* = 0.821), neuromodulation (F_3,56_ = 0.783 *P* = 0.509), nor any interaction between time and neuromodulation (F_9,127_ = 0.971, *P* = 0.467) on background EMG post-exercise. Hence, all TMS and PNS measurements were essentially taken when participants consistently held a 5% maximum EMG contraction.

### Fatigue

While there was an expected main effect of time (F_9,122_ = 18.7, *P* < 0.001), there was no main effect of neuromodulation (F_3,53_ = 0.067, *P* = 0.977) nor interaction between time and neuromodulation (F_27,379_ = 0.484, *P* = 0.987) on MVC force during fatiguing exercise (Fig. 3(A)). Across all neuromodulation conditions, when compared to the beginning of the exercise (Fatigue 1), MVC force progressively declined from the third through to the tenth contraction (*P* < 0.05) (Fig. 3(A)). Post-exercise (Fig. 3(B)), there was a main effect of time (F_3,33_ = 58.7, *P* < 0.001) on MVC force but, not neuromodulation (F_3,48_ = 0.141, *P* = 0.935). There was also an interaction between time and neuromodulation (F_9,129_ = 2.11, *P* = 0.033) on MVC force post-exercise. Force remained attenuated at 0-, 10– and 20-minutes post-exercise across all neuromodulation conditions (*P* < 0.001) (Fig. 3(B)).

**Fig. 3.**
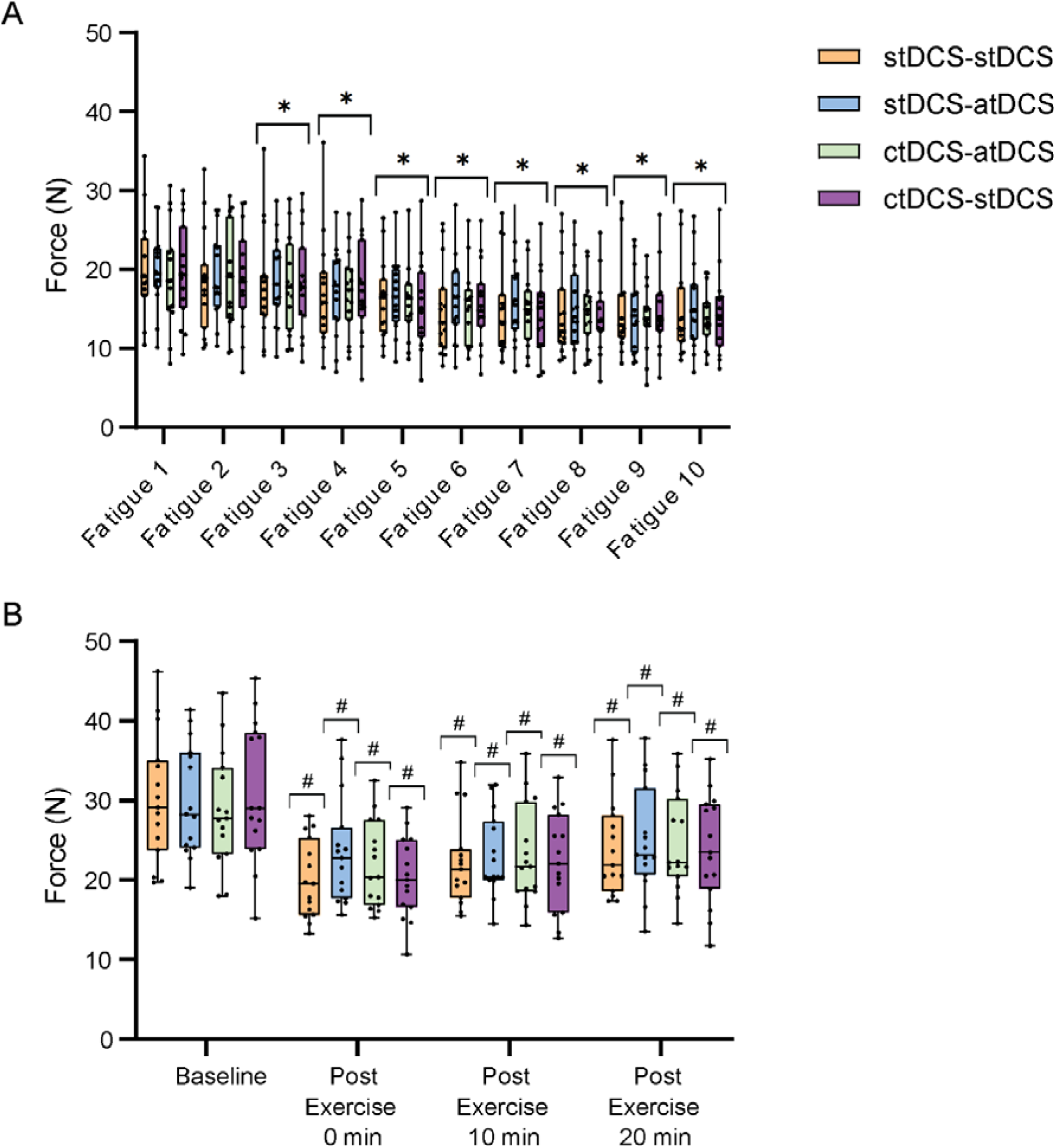
Mean MVC force during– (A) and post– (B) fatiguing exercise displayed over time and across neuromodulation conditions (stDCS-stDCS, stDCS-atDCS, ctDCS-atDCS, ctDCS-stDCS) for fifteen young healthy adults. Bonferroni’s post hoc tests: * indicates significant difference from fatiguing contraction 1 (Fatigue 1) (*P* < 0.05); # denotes significant difference from baseline (*P* < 0.05)

## DISCUSSION

### Main findings

The current study is the first to explore the interaction between metaplastic neuromodulation and neuromuscular fatigue in young, healthy individuals. Data on tDCS-induced modulation of MEP amplitudes in young healthy participants showed no significant shift in corticospinal excitability with ctDCS-atDCS compared to stDCS-atDCS, ctDCS-stDCS or excitability facilitation induced by the fatiguing exercise itself (stDCS-stDCS). Likewise, there was no benefit of tDCS, either primed or not primed, for the development of neuromuscular fatigue during exercise. Since this study revealed no significant benefit of ctDCS-atDCS on enhancing corticospinal excitability or reducing neuromuscular fatigue during exercise, the prospect of harnessing metaplasticity to ameliorate neuromuscular fatigue in young healthy individuals remains elusive.

### tDCS effects on corticospinal excitability

Our current understanding of an enhancement of MEP amplitudes during fatiguing maximal efforts (i.e. MVCs) is attributed to increases in corticospinal excitability (Taylor et al., 1996), arising as a counteractive mechanism to boost contractile force as fatigue develops (Gandevia et al., 1996). The notion that corticospinal excitability increases during single-joint fatiguing exercise (Benwell et al., 2007; Otieno et al., 2019) is verified in the present study by an enhancement of MEP amplitudes in the stDCS-stDCS condition. The Bienenstock-Cooper-Munro theory states that the threshold for synaptic modification dynamically and bi-directionally adjusts as a function of former activity (Bienenstock et al., 1982). Although it is understood that the induction of excitability modification is sensitive to the state of the network enforced by prior synaptic activity (Abraham and Bear, 1996; Abraham and Tate, 1997; Bienenstock et al., 1982; Siebner et al., 2004; Lang et al., 2004), we found no differences in MEP amplitudes between the explored neuromodulation conditions. Hence, our hypothesis of an augmentation in corticospinal excitability facilitation with ctDCS primed atDCS applied simultaneously with fatiguing exercise is not confirmed in this study.

Maximal compound muscle action potential, or Mmax, signifies full activation of the motor neurone pool of a muscle (Magladery et al., 1951). Studies reveal a decrease in Mmax amplitude in the presence of muscular fatigue (Crone et al., 1999). Therefore, in the present study, it is likely that the attenuation in Mmax is a result of the exercise protocol inducing fatigue. At a physiological level, an impairment of neuromuscular propagation is a probable cause for this attenuated Mmax (Fuglevand et al., 1993). Interestingly, in comparison to stDCS-stDCS neuromodulation, Mmax at the beginning of the fatiguing exercise (30-seconds post-test tDCS) was greater in stDCS-atDCS and ctDCS-stDCS conditions. The underlying reason for the differences in Mmax between sessions is not clear but, is likely due to individual variability in Mmax amplitude at baseline, rather than an influence of tDCS per se.

Notably, there was a lack of difference in corticospinal excitability between stDCS-atDCS and stDCS-stDCS neuromodulation in this study that conflicts with previous investigations exploring physiological and behavioural outcomes of tDCS (Cogiamanian et al., 2007; Kidgell et al., 2013; Nitsche and Paulus, 2000; Nitsche and Paulus, 2001; Nitsche et al., 2002; Williams et al., 2013). Earlier studies examining the application of atDCS during muscle activity have revealed increases in corticospinal excitability (Hendy and Kidgell, 2013; Kim and Ko, 2013). The lack of difference between stDCS-atDCS and stDCS-stDCS in the current study compared to similar research in the field (Cogiamanian et al., 2007; Williams et al., 2013) may be due to variation in methodologies and study designs. For example, unlike the present study, many tDCS investigations have failed to blind the experimenter involved in the research (Cappa, 2008; Marangolo et al., 2011; Monti et al., 2008; Norise et al., 2017; Shah-Basak et al., 2015); increasing the risk of bias. Additionally, the experimental parameters of tDCS vary among studies. Factors such as timing, intensity, and duration of tDCS application are known to influence tDCS outcomes (Sellaro et al., 2016). In this study, tDCS was applied at an intensity of 1 mA yet, other studies in the area commonly employ a tDCS intensity of 1.5 – 2 mA (Abdelmoula et al., 2019; Cogiamanian et al., 2007; Fujiyama et al., 2017; Kan et al., 2013). It has been established, however, that higher tDCS intensities does not necessarily increase corticospinal excitability if more than 1.5 mA is administered (Kan et al., 2013). Likewise, the duration of tDCS application in this study was 15 minutes of priming and 12 minutes during the exercise task but, studies reporting an effect of stDCS-atDCS apply tDCS during performance of a motor task for 20 minutes (Christova et al., 2015; Fujiyama et al., 2017). Batsikadze and colleagues have demonstrated that an increase in tDCS intensity or duration is not strictly associated with an enhancement of its efficacy but may in fact alter the direction of its effects (Batsikadze et al., 2013). Inter-individual variability of response to tDCS may also contribute to the conflicting outcomes of the present study to that of prior investigations (Chew et al., 2015; Xian et al., 2023). In fact, it has been observed that the after-effects of neuromodulation are variable within and between participants in relation to direction, duration, and magnitude (Huang et al., 2017) and influenced by electrode and skull characteristics (Antonenko et al., 2021). A variety of factors may play a role in the neuromodulation response variability including sex, attention, physical activity levels and optimal stimulation dose (Fertonani and Miniussi, 2017; Guerra et al., 2020; Huang et al., 2017; Li et al., 2015; López-Alonso et al., 2014; Ridding and Ziemann, 2010; Rudroff et al., 2020). Hence, future studies exploring the relationship between corticospinal excitability and different tDCS parameters are necessary to decipher the most appropriate arrangement for enhancing tDCS outcomes (Amann et al., 2022).

### tDCS effects on fatigue

The ability of neuromodulation to mutually modify motor performance and corticospinal excitability highlights the possibility of exploiting neuromodulation to influence mechanisms of fatigue (Christova et al., 2015; Fujiyama et al., 2017). The present study reveals an attenuation of MVC force during exercise regardless of the neuromodulation condition. MVC remained significantly lower than baseline across all four conditions at 20 minutes post-exercise. Kan et al. (2013) reported similar results in their study, wherein atDCS had no effect on MVC strength or time-to-task-failure (TTF) of the elbow flexors (Kan et al., 2013). However, unlike the delivery of tDCS during exercise in the present study, Kan et al. (2013) applied atDCS prior to exercise. Conversely, Williams et al. (2013) reached differing conclusions in their study, whereby application of atDCS during fatiguing exercise led to an improvement in performance. A fundamental distinction in the protocol that could justify the contradictory result concerns the quantification of fatigue. Williams et al. (2013) assessed fatigue by TTF in submaximal isometric contractions (as opposed to intermittent MVCs in the current study). Consequently, it is probable that amelioration of fatigue via tDCS is task dependent and regulated, at least partially, by exercise intensity.

Since development of fatigue is related to an increase in corticospinal excitability (Benwell et al., 2007; Otieno et al., 2019), a parallel between fatigability and excitability of outcomes is expected. Earlier research has demonstrated an attenuation of fatigue upon delivery of atDCS concurrently with exercise (Oki et al., 2016; Williams et al., 2013). Yet, investigations on priming ctDCS prior to concurrent application of atDCS during a task have illustrated enhanced corticospinal excitability and motor performance when compared to atDCS with no priming (stDCS) (Christova et al., 2015; Fujiyama et al., 2017). Hence, we anticipated that excitability modulation via priming ctDCS prior to atDCS delivered simultaneously during fatiguing exercise would attenuate fatigue compared to atDCS primed by stDCS. However, onset of fatigue and lack of recovery in force post-exercise was consistent across all investigated tDCS paradigms as no differential tDCS modulation of corticospinal excitability was present in this study.

### Limitations

The current study has some considerations that should be addressed. Firstly, it could be disputed that capitalisation of metaplasticity was not effective as the priming alone caused no modification in MEP. Yet, even when priming stimulation fails to produce noticeable variations in excitability, subsequent synaptic alterations have been shown to occur (Abraham, 2008; Abraham and Bear, 1996; Sidhu, 2021). Moreover, several variables are well-established in influencing the outcomes of tDCS. Although we are aware of hormones and the potent regulatory role they play in plasticity (Abraham, 2008; Ansdell et al., 2019), we did not control for the menstrual cycle in female participants and is considered a limitation. Additionally, sex differences in fatigability are documented (Ansdell et al., 2020; Hunter, 2014). Females are commonly described as having greater resistance to fatigue because of greater availability of oxygen during exercise (Ansdell et al., 2019; Ansdell et al., 2020). Regarding study techniques, neuronavigation was not employed in our study. Neuronavigation systems allow for accurate identification of the brain area of interest by utilisation of an individual’s magnetic resonance imaging data (Sparing et al., 2010). While neuronavigation is common in TMS studies for precise positioning of the magnetic coil, it is of greater importance in studies exploring brain regions other than M1 (e.g., frontal cortex) (Herwig et al., 2001; Sparing et al., 2010). It could also be argued that the relatively homogenous young healthy study sample is a limitation. However, investigating the basic neurophysiological mechanisms of fatigue in a healthy cohort has broader significance for the field. Lastly, there is some indication of increased variability if fewer than 20 simultaneous MEP responses are averaged (Biabani et al., 2018). Thus, the 5 TMS-evoked MEP responses during fatiguing exercise, and 15 at every other time point, should be acknowledged.

### Conclusion

This work offers a novel application of tDCS to modulate corticospinal excitability and fatigability in young, healthy adults. In contrast to prior investigations, we observed no differential modulation of corticospinal excitability or fatigue with delivery of atDCS during isometric single-joint fatiguing exercise. Consequently, priming ctDCS prior to fatiguing exercise combined with atDCS failed to augment corticospinal excitability facilitation or attenuate fatigue. Overall, the present results highlight some limitations in the use of tDCS to attenuate muscle fatigability. Further exploration to establish the efficacy of tDCS and the underlying neurophysiological mechanisms in mitigating neuromuscular fatigue in both normal and pathological settings is warranted.

## STATEMENTS AND DECLARATIONS

### Ethical approval

This study was approved by the Human Research Ethics Committee at the University of Adelaide and performed in accordance with the ethical standards as laid down in the 1964 Declaration of Helsinki and its later amendments.

### Consent to participate

Written informed consent was obtained from all participants before involvement in the study.

## Abbreviations

AMT: active motor threshold
atDCS: anodal transcranial direct current stimulation
ctDCS: cathodal transcranial direct current stimulation
EMG: electromyography
FDI: first dorsal interosseous
LMM: linear mixed model
LTD: long term depression
LTP: long term potentiation
M1: primary motor cortex
MEP: motor evoked potential
Mmax: maximum M-wave
MVC: maximal voluntary contraction
PNS: peripheral nerve stimulation
stDCS: sham transcranial direct current stimulation
tDCS: transcranial direct current stimulation
TMS: transcranial magnetic stimulation
TTF: time-to-task-failure

## Acknowledgements

This work was supported by the Adelaide Medical School Honours Scholarship awarded to MRB.

## Glossary

Corticospinal Excitability: the excitability of the pathway from the cortical site of neuronal depolarization to spinal motoneuron depolarization.
Long Term Depression (LTD): a phenomenon where the cumulative activation of inputs to specific neural pathways produces a decrease in the excitability of these neurons.
Long Term Potentiation (LTP): a phenomenon where repeated electrical stimulation of inputs to specific neural pathways produces an increase in the excitability of these neurons.
Motor Evoked Potential (MEP): the electrical signals recorded from the descending motor pathways or from muscles following stimulation of motor pathways within the brain.
Neuromodulation: the alteration of neuronal and synaptic properties by neurons themselves, substances released by neurons, or by technology that acts directly upon nerves.
Neuromuscular Fatigue: a multifactorial phenomenon that can occur in various sites along the pathway of force production, causing a reduced ability to generate a desired muscle force.
Neuroplasticity: also known as neural plasticity or brain plasticity, is a process that involves adaptive structural and functional changes to the brain.
Trancranial Direct Current Stimulation (tDCS): a non-invasive form of neuromodulation used to modulate corticospinal excitability by use of constant, low, direct current delivered via electrodes on the head.
Transcranial magnetic stimulation (TMS): a non-invasive form of brain stimulation used to test and modify corticospinal excitability via electromagnetic induction.

